# Role of Adenosine Deaminase in Prostate Cancer Progression

**DOI:** 10.1101/2023.08.09.552704

**Authors:** Christy Charles, Stacy M. Lloyd, Danthasinghe Waduge Badrajee Piyarathna, Jie Gohlke, Uttam Rasaily, Vasanta Putluri, Brian W. Simons, Alexander Zaslavsky, Srinivas Nallandhighal, Nallasivam Palanisamy, Nora Navone, Jeffrey A. Jones, Michael M. Ittmann, Nagireddy Putluri, David R. Rowley, Simpa S. Salami, Ganesh S. Palapattu, Arun Sreekumar

## Abstract

Prostate cancer (PCa) is the second most common cancer and constitutes about 14.7% of total cancer cases. PCa is highly prevalent and more aggressive in African American (AA) men when compared to European-American (EA) men. PCa tends to be a highly heterogeneous malignancy with a complex biology that is not fully understood. We use metabolomics as a tool to understand the mechanisms behind PCa progression and disparities in its clinical outcome. A key enzyme in the purine metabolic pathway, Adenosine deaminase (ADA) was found upregulated in PCa. ADA was also associated with higher-grade PCa and poor disease-free survival. The inosine-to-adenosine ratio which is a surrogate for ADA activity was high in the urine of PCa patients and higher in AA PCa compared to EA PCa. To understand the significance of high ADA in PCa, we established ADA overexpression models and performed various in vitro and in vivo studies. Our studies have revealed that an acute increase in the expression of ADA during later stages of tumor development enhances in vivo growth in multiple pre-clinical models. Further analysis reveals that this tumor growth could be driven by the activation of mTOR signaling. Chronic ADA overexpression shows alterations in the cells’ adhesion machinery and a decrease in the adhesion potential of the cells to the extracellular matrix in vitro. Loss of cell-matrix interaction is critical for metastatic dissemination, suggestive of ADA’s role in promoting metastasis. This is consistent with the association of higher ADA expression with higher-grade tumors and poor patient survival. Overall, our findings suggest that increased ADA expression may promote PCa progression, specifically tumor growth and metastatic dissemination.

---

Prostate cancer (PCa) is the most commonly diagnosed cancer in American men. One in eight men are diagnosed with PCa in their lifetime. And ∼300,000 men are estimated to have PCa in 2023. Currently, about 3.1 million men are living with the disease in the United States[1]. There is a striking racial disparity that exists within PCa. African American (AA) men experience higher incidence and mortality rates compared to European American (EA) men [2]. Prostate tumors have a complex biology, with both aggressive and non-aggressive tumors being highly differential. The heterogeneity of the tumors is not fully understood and there are more knowledge gaps on the tumor characteristics and progression.

Our group focuses on identifying the metabolic pathways and the associated molecular mechanisms driving PCa progression. Our goal is to identify the metabolic alterations associated with PCa and understand its function in the tumor setting. Previously, we studied the metabolic landscape of PCa patients and reported alterations within the methionine-homocysteine pathway in AA PCa [3]. Methionine is an essential amino acid, and its metabolism leads to homocysteine/cysteine and adenosine/inosine production. In the current study, we examine the enzyme adenosine deaminase (ADA, EC 3.5.4.4.), that regulates the adenosine-inosine axis.

ADA catalyzes the irreversible conversion of adenosine to inosine. Biologically, the well-known significance of ADA activity is that the inosine generated by the enzyme counteracts the immunosuppressive effects of adenosine. ADA is well-studied in the context of the immune system and immunodeficiencies, such as Severe Combined Immunodeficiency (SCID). In immunodeficient diseases, ADA levels are lower, leading to the build-up of adenosine, which causes severe DNA damage and lymphocytotoxicity [4]. ADA, however, is elevated in certain types of cancers and is associated with poor survival [5]. ADA inhibition was shown to prevent breast cancer progression in mice and induce cytotoxicity in cervical cancer and malignant pleural mesothelioma cells [6–8]. However, the biological effects of ADA in any solid tumor have not been fully characterized. In the current study, we sought to elucidate the effects of ADA on tumor progression in PCa. Our goal is to delineate the intrinsic tumor effects of ADA in the context of PCa development and progression. As a result, we have assessed the clinical levels of ADA in PCa and compared the levels of expression across Gleason grades, as well as between AA and EA PCa patients. We have also extensively studied the effects of ADA elevation in PCa progression using in vitro and in vivo immunocompromised murine models. We found that elevated ADA in PCa supports tumor growth and aggressive behavior.

## Methods

### Clinical samples

All clinical samples used in this study were obtained using informed consent and the approval of the Institutional Review Board at Baylor College of Medicine (Protocol title: Integrative metabolomics of cancer progression; Protocol number: H-28445; Expiration date: 09/20/2026) and collaborating institutions, including the National Cancer Institute (Study PI: Dr. Stefan Ambs). The urine samples were kindly provided by Dr.Stefan Ambs (National Cancer Institute). Tissue microarrays for the RNA *in situ* hybridization study were obtained from Henry Ford Health. Tissue microarrays for ADA immunostaining were obtained from the Baylor College of Medicine Human Tissue Acquisition and Pathology Core. The urine samples were obtained from the NCI-Maryland Prostate Cancer case-control study. The clinical characteristics of the urine samples are described in Table 1.

**Table 1.**
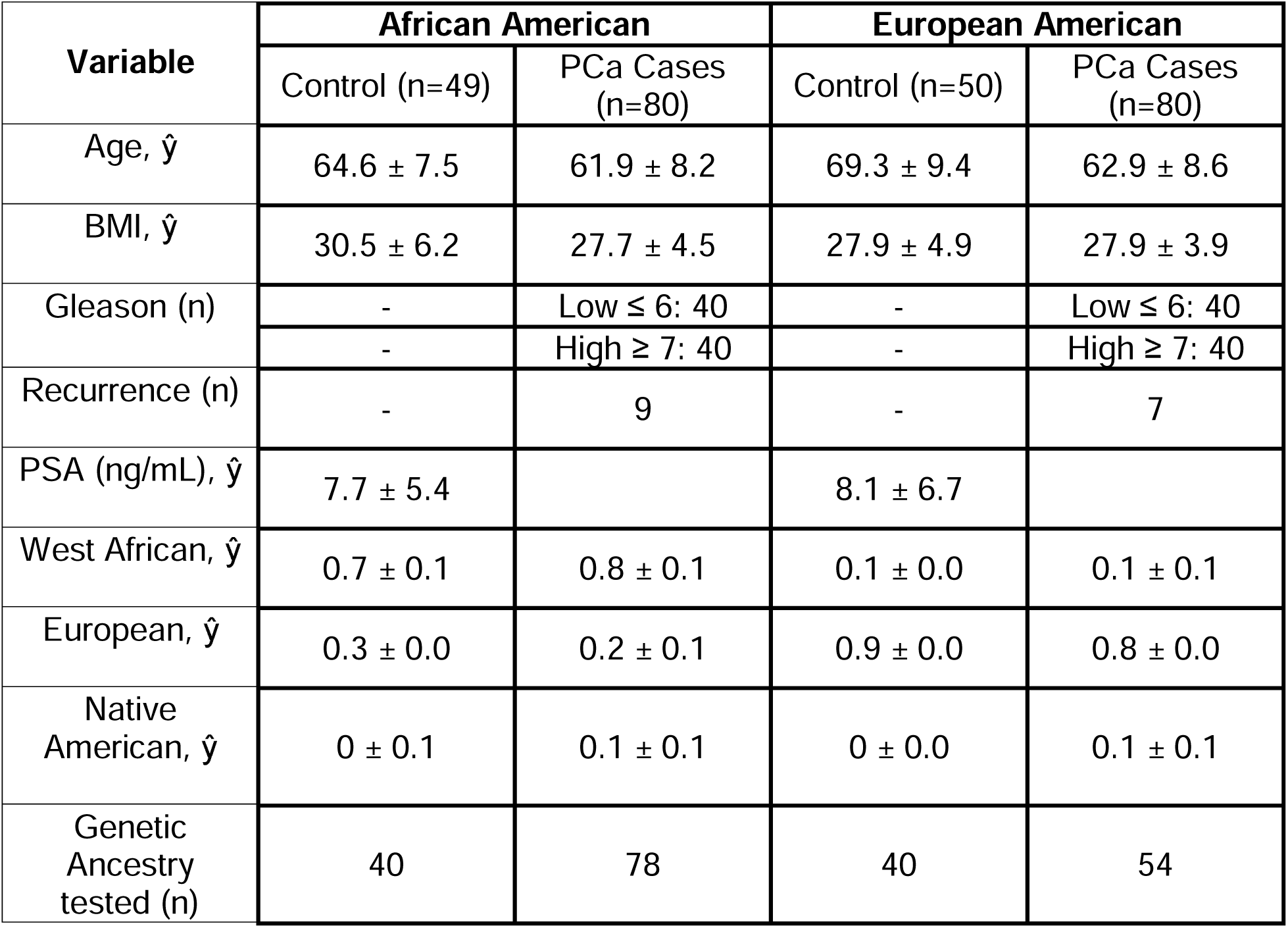
Summary of clinical and ancestry data for PCa case-control urine cohort.

| Variable | African American |  | European American |  |
| --- | --- | --- | --- | --- |
|  | Control (n=49) | PCa Cases (n=80) | Control (n=50) | PCa Cases (n=80) |
| Age, $\hat{y}$ | 64.6 ± 7.5 | 61.9 ± 8.2 | 69.3 ± 9.4 | 62.9 ± 8.6 |
| BMI, $\hat{y}$ | 30.5 ± 6.2 | 27.7 ± 4.5 | 27.9 ± 4.9 | 27.9 ± 3.9 |
| Gleason (n) | - | Low ≤ 6: 40 | - | Low ≤ 6: 40 |
| | - | High ≥ 7: 40 | - | High ≥ 7: 40 |
| Recurrence (n) | - | 9 | - | 7 |
| PSA (ng/mL), $\hat{y}$ | 7.7 ± 5.4 | | 8.1 ± 6.7 | |
| West African, $\hat{y}$ | 0.7 ± 0.1 | 0.8 ± 0.1 | 0.1 ± 0.0 | 0.1 ± 0.1 |
| European, $\hat{y}$ | 0.3 ± 0.0 | 0.2 ± 0.1 | 0.9 ± 0.0 | 0.8 ± 0.0 |
| Native American, $\hat{y}$ | 0 ± 0.1 | 0.1 ± 0.1 | 0 ± 0.0 | 0.1 ± 0.1 |
| Genetic Ancestry tested (n) | 40 | 78 | 40 | 54 |

### Fluorescent RNA *in situ* hybridization

Slides were incubated at 60°C for 1 hour. Deparaffinization was done by immersing in xylene twice for 5 minutes each with periodic agitation. The slides were then immersed in 100% ethanol twice for 3 minutes each with periodic agitation, then dried at 60° C for 5 minutes. Tissues were treated with 5-8 drops of H_2_O_2_ for 10 minutes at room temperature. Slides were washed twice in distilled water and then boiled in 1X Target Retrieval for 15 minutes. Slides were washed twice in distilled water, immersed in 100% EtOH for 3 minutes, and then dried for 5 minutes at 60°C. Tissues were marked using a hydrophobic barrier pen (cat# H-4000 Vector Labs, Burlingame, CA), allowed to dry, and then treated with Protease Plus for 30 minutes at 40°C in a HybEZ Oven (cat# 310010, Advanced Cell Diagnostics, Hayward, CA). Slides were washed twice in distilled water and then treated with the ADA probe (cat# 490141, Advanced Cell Diagnostics, Hayward, CA) for 2 hours at 40°C in the HybEZ Oven. Slides were then washed in 1X Wash Buffer (cat# 310091, Advanced Cell Diagnostics, Hayward, CA) twice for 2 minutes each. All slides were then treated with Amp 1 for 30 minutes, Amp 2 for 30 minutes, and Amp 3 for 15 minutes, all at 40°C in the HybEZ oven with two washes in 1X Wash Buffer for 2 minutes each after each step. The slides were treated with HRP-C1 for 15 minutes, Opal 520 (cat# FP487001KT, Akoya Biosciences) diluted 1:1500 in TSA Buffer for 30 minutes, HRP Blocker for 15 minutes, and HRP-C3 for 15 minutes all at 40L C in the HybEZ oven with two washes in 1X Wash Buffer for 2 minutes each after each step. DAPI was added to all slides for 30 seconds at room temperature, which was then rinsed off. 1-2 drops of ProLong Gold Antifade Mountant (cat# P36930, Molecular Probes, Eugene, OR) were added, and slides were covered with coverslips. H_2_O_2,_ Target Retrieval, Protease Plus, Wash Buffer, Amps 1-3, HRP-C1-3, TSA Buffer, HRP Blocker, and DAPI are all part of the RNAscope Multiplex Fluorescent Reagent Kit v2 (cat# 323100, Advanced Cell Diagnostics, Hayward, CA).

### Immunohistochemistry

Slides were deparaffinized by three xylene washes and rehydrated with a graduated alcohol series. Antigen retrieval was performed at 60°C in citrate-based antigen unmasking solution diluted 1:100 (Vector Labs, Burlingame, CA) for 20 minutes. Slides were washed with 1X Phosphate Buffered Saline-Tween (PBS-T) and treated with BLOXALL endogenous blocking solution for 10 minutes and blocked with 2.5% Normal horse serum (Vector Labs, Burlingame, CA) for 20 minutes. Primary antibody incubation was performed overnight at 4°C with 0.5 mg/mL Mouse ADA antibody (MBS1752027, MyBioSource, San Diego, CA). The secondary antibody and the substrate for the detection are added from the Vectastain Universal Elite ABC Immunohistochemistry Kit (Cat # PK-8200, Vector Labs, Burlingame, CA). Slides were incubated in secondary antibody for 20 minutes and washed with PBS-T. The substrate was added after the wash. Slides were briefly rinsed in water before counterstaining with Gill’s No. 2 Hematoxylin (Cat # 10143746, VWR, Radnor, PA) for 1 min. Slides were rinsed in water and incubated with Scott’s Bluing Reagent (Cat # RC669732, VWR, Radnor, PA) for 1 min. Slides were washed and dehydrated in a graduated alcohol series followed by xylene dips before mounting. The specificity of the ADA antibody was verified by using control slides in the study. The control slides were generated from xenografts formed in mice using control and ADA-overexpressing prostate cell lines. Colon tissues were also used as an additional positive control. Additionally, the antibody was also verified using a western blot. The staining was scored by an experienced genitourinary pathologist in the core lab, who assigned each section an intensity and extent score. These two values were multiplied (intensity x extent) to produce a final score for each core sample.

### Cell Lines

The MDA-PCa-2a cell line was a gift from Dr. Nora Navone (MD Anderson Cancer Center). LNCaP was obtained from the Tissue Culture Core at Baylor College of Medicine. The 22Rv1 cell line was obtained from ATCC (CRL-2505). C4-2B was a gift from Dr. Nancy Weigel (Baylor College of Medicine). Stromal line HPS-19I was a gift from Dr. David Rowley (Baylor College of Medicine). Short Tandem Repeat (STR) analysis was completed for all cell lines to verify their authenticity at the MD Anderson Cytogenetics and Cell Authentication Core. Mycoplasma tests (MycoAlert^TM^ from Lonza, Anaheim, CA) were conducted routinely to ensure the cells remained free of mycoplasma contamination.

### Cell Culture

MDA-PCa-2a was cultured in BRFF-HPC1 media (AthenaES, Baltimore, MD) with 20% fetal bovine serum (FBS, Hyclone Labs, Thermo Scientific, Rockford, IL) and 0.1% penicillin-streptomycin (Hyclone Labs, Thermo Scientific, Rockford, IL). MDA-PCa-2a cells were grown in plates/flasks coated with a fibronectin coating mix (AthenaES, Baltimore, MD). LNCaP, 22Rv1, and C4-2B were grown in RPMI-1640 media (Invitrogen Corp., Carlsbad, CA) supplemented with 20% fetal bovine serum and 0.1% penicillin-streptomycin. The cells were maintained at 37°C, 5% CO_2,_ and 95% humidity. The cells were passaged or harvested when they reached about 80% confluency. The cells were treated with 0.25% Trypsin-EDTA (Gibco, Life Technologies, Grand Island, NY) for about 5-10 minutes. Once all the cells were off the substratum, culture media was added for neutralization. The cells were then counted using Trypan blue stain (Life Technologies, Eugene, OR) and Countess^TM^ Automated Cell Counter (Life Technologies, Singapore). The cells were seeded at appropriate densities according to the culture dishes used or the experimental requirements.

### Lentiviral Transduction

To generate a stable ADA overexpressing system, the cells were transduced with lentiviral cDNA open reading frames (Precision LentiORFs, Horizon Discovery, Cambridge, UK). The precision lentiORF for ADA (Catalog #OHS5836-EG100, Clone: OHS5899-202616178 (PLOHS_100005311)) and a non-target ORF (Catalog #OHS5833) was used as a control. A multiplicity of infection (MOI) of 5 was used for lentiviral transduction. About 8 µg/mL of blasticidin (Gibco, Life Technologies, Grand Island, NY) was for selection. Additionally, the cells were also checked for the presence of GFP (ADA overexpression) and RFP (control) under the fluorescence microscope to verify transduction efficiency. For creating a knockdown rescue in the ADA OE cells, transduction with shRNA lentivectors (Gentarget Inc, San Diego, CA) was done. shRNA targeting ADA and non-target shRNA (vector control) were used (catalog #LVS-1002). An MOI of 5 was used for lentiviral transduction, and 1 µg/mL puromycin was used for the selection of effectively transduced cells. The inducible ADA overexpression lines were developed at the Cell-Based Assay Screening Service (CBASS) at Baylor College of Medicine. The lines were generated by transduction of tet-inducible lentiviral vector for ORF expression (Addgene, pInducer 20, catalog # 44012). The ADA gene was cloned into the overexpression cell line and the 11^th^ beta-strand of GFP was used as a doxycycline control (Dox-control).

### Quantitative Real-Time Polymerase Chain Reaction

RNA extraction from cells was performed using the Aurum total RNA Mini Kit from Biorad. For tissues, RNA was extracted using the RNeasy mini kit (Qiagen, Hilden, Germany). Extracted RNA was quantified and its purity (absorbance at 260/280 nm) was verified using a spectrophotometric plate reader (Synergy HTX multi-mode reader, Biotek Instruments, Winooski, VT). RNA was reverse transcribed into cDNA using the cDNA Superscript Mix (Quanta Biosciences, cat#95048-500) and RT-qPCR was done using SYBR green (Life Technologies, cat#4385614). 18S or RPL30 were used as appropriate housekeeping controls. Primers used in this study are listed in Table 2.

**Table 2.**
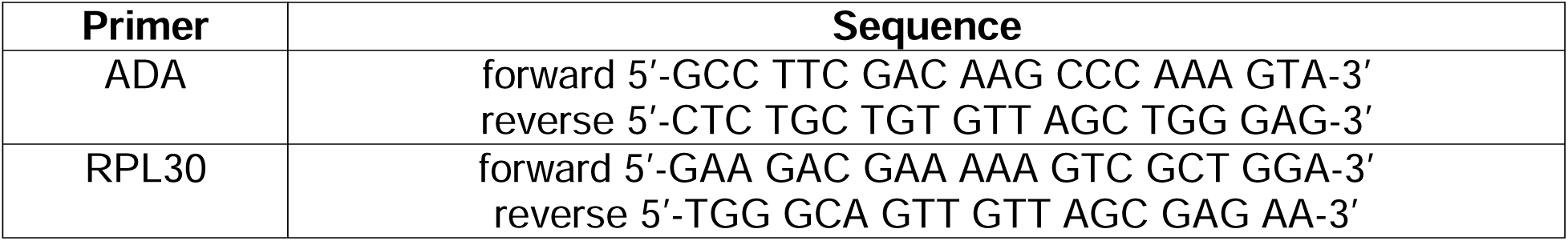
Primers used in this study.

### Protein extraction and estimation

Cells were washed in ice-cold PBS thrice to remove residual culture media. RIPA buffer was added in appropriate amounts to the cells and incubated for 10 minutes in ice. The cells were then sonicated and spun down. The supernatant (cell lysate) was collected, and the total protein was quantified using Pierce^TM^ BCA Protein Assay Kit (Thermo Fisher, Waltham, MA).

### Western Blot

Cell lysates with 30 µg protein were mixed with 2x Laemmli sample buffer (Biorad, catalog #1610737) in a 1:1 ratio and heated at 95°C for 10 minutes. Then the prepared samples were loaded onto the lanes of a 4-15% Mini-PROTEAN Tris PAGE gel (Bio-rad #4561086). Electrophoresis was carried out at 100 V for about 1.5-2 hours. The separated proteins were transferred from the gel to the polyvinyl difluoride (PVDF) membrane (Immobilon-P, 0.45 µm, Merck Millipore Ltd, Ireland) at 200 mA for 1.5 hours. The transferred blot was blocked using 5% skimmed milk and incubated with the ADA antibody (MyBioSource, #MBS1752027) overnight. The blot was washed with TBST for 10 minutes three times and then incubated with the appropriate secondary antibody for 1 hour. The blot was again washed with TBST for 10 minutes thrice. Finally, the blots were probed using a chemiluminescent substrate (Thermo Fisher, cat# 34579) in the Chemi-Doc Imaging system (Biorad, Hercules, CA). Polyclonal rabbit α/β-tubulin antibody (CST, #2148S) was used as an internal control.

### ADA Enzyme assay

ADA assay kit (DZ117A; Diazyme Laboratories, Inc., Poway, CA) was used for the determination of ADA activity in cell lysates. About 4ug of protein was used for the assay. The manufacturer’s protocol was followed to assess the ADA enzyme activity.

### LC/MS

Adenosine and inosine levels in urine were measured by LC/MS. Adenosine (CNLM-3806-CA-PK, Cambridge Isotope Laboratories, Inc., Andover, MA) and inosine (NLM-4264-0.01, Cambridge Isotope Laboratories, Inc., Andover, MA) standards for concentrations 5-100 ng/mL were prepared in a urine matrix (MSG5000, Golden West Diagnostics, Temecula, CA). 10 µL of urine was mixed with 90 µL of methanol: water (1:1) and internal standards (Labeled tryptophan (CNLM-2475-H-PK, Cambridge Isotope Laboratories, Inc., Andover, MA), zeatine and inosine) were spiked into this. Samples were then loaded into LC/MS sampler vials and injected into the mass spectrometer.

To evaluate the adenosine and inosine levels in cell lines, the cells were harvested and washed thrice to remove any residual media. About 100 µl water and 5 µl internal standard (Labeled adenosine and inosine) were added to the cells. The internal standards were dissolved in a 1:1 methanol-water mix by vortexing for 5 min. The cells were homogenized by probe-sonication for 30 seconds (in ice). 50 µl DTT (500 mM-freshly prepared) was added, and samples were vortexed for 20 seconds. The samples were then incubated at 65 ^□^C for 30 min at 300 rpm. After incubation, the samples are kept on ice for a minute. 400 µl cold methanol was added and mixed well by vortexing for 5 min. This mixture was centrifuged, and the supernatant was collected. The collected supernatant was then dried. After drying, 50 µl of a 1:1 methanol-water was added. To ensure proper dissolution, samples were vortexed for 5 min and sonicated for 5 min. The samples were then centrifuged for 5 min. Finally, samples were transferred into the inserts of LC/MS vials and injected into the mass spectrometer. The chromatography platform used to separate all the metabolites in the study was Agilent Zorbax SB-CN (3×100mm;1.8μm) column; mobile phase, A: 0.1% formic acid in HPLC grade water, B: 0.1% formic acid in HPLC grade acetonitrile. Acetonitrile, methanol, and water for high-performance liquid chromatography (HPLC) were purchased from Burdick & Jackson (Morristown, NJ).

### Adhesion assay

The cell-ECM/surface adhesion was measured using the Agilent xCELLigence Real-Time Cell Analysis (RTCA) DP (dual purpose) instrument. About 10,000 cells are seeded onto Cell invasion and migration (CIM) -Plates (G082975-Agilent, San Diego, CA). Adhesion was automatically measured by the instrument at regular time intervals for 100 hours.

### In vivo experiments

Male NOD-SCID-Gamma (NSG) mice were obtained from Baylor College of Medicine, and male athymic nude mice (Strain #553) were purchased from Charles River (Frederick, MD) for these studies. The experiments started when the mice were between 4-6 weeks old. Nude mice were used to grow LNCaP tumors. All other cell line xenograft studies were done in NSG mice. These mice were housed in germ-free cages with a maximum of 4 mice per cage in the immunodeficient mice facility. Mice were randomized into experimental groups based on the cage. The sample size for in vivo studies was determined based on statistical power calculations and past experiments from our laboratory. All in vivo experiments were started with 10 mice per group. Animals lost to attrition were excluded from the study. And data from the remaining mice were used for evaluation. The studies were not blinded, and outliers were not excluded. The experiments were carried out and the health of the mice was monitored per the animal protocol approved by the Institutional Animal Care and Use Committee (IACUC) of Baylor College of Medicine (Protocol title: Integrative metabolomics of cancer detection and progression; Protocol number: AN-5676; Expiration date: 02/13/2026).

### In vivo: tumor take and growth studies

To evaluate the tumor take and growth, cells were injected subcutaneously. 5 million cells were used for MDA-PCa-2a and C4-2B and 3 million cells for 22Rv1. For LNCaP tumors, 500,000 19I stromal cells were added along with 5 million LNCaP cells. All the cells were mixed with 50µL Matrigel and 50µL culture media. 100µL of the cells-matrigel mixture was injected per mouse. Injections were done after the mice were anesthetized using 2% vaporized isoflurane.

Two tumor growth studies were done: 1) ADA was overexpressed constitutively 2)ADA levels were elevated by doxycycline induction after tumorigenesis. For the first study, the constitutive ADA OE lines from in vitro studies (MDA-PCa-2a and LNCaP) and an ADA-inducible cell line (22Rv1) kept under constant induction were used. The 22Rv1 ADA OE cell line used for this experiment is an inducible cell line, placed under constant ADA induction in culture, during injection, and tumor growth via continuous doxycycline exposure. The corresponding control used is a doxycycline control, which is also maintained under constant doxycycline exposure just like the ADA OE line. This eliminates any doxycycline-mediated consequences.

For the second study, to induce ADA post-tumor take doxycycline was added to the drinking water at the concentration of 0.5mg/mL after tumors formed (∼50 mm^3^). The water was changed twice a week and a special care instructions form was placed to indicate deviation from regular drinking water. Two controls were used in this study, an uninduced control without doxycycline exposure and a doxycycline control (dox-control, where GFP is induced instead of ADA)

The tumor dimensions were measured using a vernier caliper once in 2 days. The volume was calculated using the formula ½ (length x breadth x width). The tumors were resected when the tumors reached 500mm^3^. The resected tumors were cut into two halves: one half was flash-frozen in liquid nitrogen and the other was fixed using 10% formalin and later paraffin-embedded (FFPE). The frozen and FFPE tissues were used for further molecular analyses.

### Reverse Phase Protein Array (RPPA)

RPPA analysis was done by the RPPA core at Baylor College of Medicine. Cell lysates were arranged on nitrocellulose-coated slides (Grace Bio-labs, Bend, OR) using Aushon 2470 Arrayer (Aushon BioSystems, Billerica, MA) and printed in three replicates (technical replicate). Three biological replicates were used for the study and each replicated was printed in triplicate. Immunolabeling was done on an automated slide stainer Autolink 48 (Dako, Carpinteria, CA) according to the manufacturer’s protocol (Autostainer catalyzed signal amplification (CSA) kit, Dako, Carpinteria, CA). Each slide was incubated with a single primary antibody at room temperature for 30 min followed by a goat anti-rabbit or mouse IgG secondary antibody. For negative control, a slide was incubated with antibody diluent instead of primary antibody. The Catalyzed Signal Amplification System kit (Dako Cytomation, Carpinteria, CA, USA) and fluorescent IRDye 680 Streptavidin (LI-COR, Lincoln, Nebraska, USA) were used for detection. The total protein for each printed spot was evaluated by staining one slide for every 20 slides with Sypro Ruby Blot Stain (Molecular Probes, Eugene, OR) according to the manufacturer’s directions. Slides were scanned using a GenePix AL4200 scanner (at 635 nm wavelength for antibody slides or 535 nm wavelength Sypro Ruby Blot-Stained slides), and the images were analyzed by GenePix Pro 7.2 software (Molecular Devices, Sunnyvale, CA). The fluorescence signal for each spot was estimated from the fluorescence intensity after subtracting the corresponding slide background signal.

To normalize the RPPA data the following method was followed. Each spot on the array has a background-subtracted foreground signal intensity (SI). If the background intensity is higher than the foreground intensity, the spot’s SI is set to a very small intensity value, 1. To normalize a sample/spot’s SI for a specific antibody, the antibody SI of each spot is subtracted by the corresponding SI of negative control and then normalized to the corresponding SI of total protein within the same group. The normalized antibody SI is calculated using the following formula,

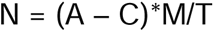

where N is the normalized antibody SI, A is the antibody SI, C is the negative control SI, M is the median SI of the spots from the same group, and T is the SI of total protein.

The differentially expressed proteins were detected by t-test at nominal P < 0.05 and FDR < 0.25. The FDR was corrected by Benjamini & Hochberg Procedure (BH method) in MDA-PCa-2a and LNCaP data. Using the differential proteins, hypergeometric enrichment analysis was conducted to identify key pathways enriched in Hallmark and KEGG (Kyoto Encyclopedia of Genes and Genomes) pathways collections) Gene sets from the Molecular Signatures Database (MSigDB). A nominal P < 0.01 and an FDR < 0.05 were used as thresholds for determining the significance.

### RNA Sequencing

RNA was extracted using the RNeasy mini kit (Qiagen, Hilden, Germany). Extracted RNA was quantified and its purity (absorbance at 260/280 nm) was verified using a spectrophotometric plate reader (Synergy HTX multi-mode reader, Biotek Instruments, Winooski, VT). Library preparation and sequencing were done at the University of Michigan Advanced Genomics Core. Library preparation and selection of poly-adenylated transcripts were done using NEBNext Poly(A) mRNA Magnetic Isolation Module (NEB, E7490) and xGen Broad-range RNA Library Prep (IDT, 1009813) with xGen Normalase UDI Primers (IDT, various). This underwent 151bp paired-end sequencing according to the manufacturer’s protocol (Illumina NovaSeq). De-multiplexed Fastq files were generated using BCL Convert Conversion Software v4.0 (Illumina). Snakemake[9] was used to manage the bioinformatics workflow and ensure reproducibility.

Quality Control (QC) and sequence alignment: The reads were trimmed using Cutadapt v2.3 [10]. The reads were evaluated with FastQC (https://www.bioinformatics.babraham.ac.uk/projects/fastqc/) (v0.11.8) to assess the quality of the data. Reads were mapped to the reference genome GRCh38 (ENSEMBL 109), using STAR v2.7.8a, and count estimates were assigned to the genes with RSEM v1.3.3 [11]. ENCODE standards were followed for alignment options in RNA-seq [12]. QC metrics from several different steps in the pipeline were combined by multiQC v1.7 [13]. After filtering low-count reads, 15328 protein-coding genes were used for differential expression analysis. The differential expression was analyzed with DESeq2 [14].

To characterize biologically significant changes in molecular signaling pathways among ADA OE and control tumors, we employed GSEA [15] to identify significantly enriched concepts in both 22RV1 and C4-2B data. In GSEA, the cumulative distribution function was constructed by performing 1000 random gene set assignments(permutations)(GSEA pre-ranked method). A nominal P < 0.05 and a FDR < 0.25 were used to threshold the concepts. Here we focused on well-defined, large-scale biological processes, termed the Hallmark (H) and KEGG (C2, Kyoto Encyclopedia of Genes and Genomes) Kyoto Encyclopedia of Genes and Genomes) Gene sets from the Molecular Signatures Database (MSigDB). Initial Principal component analysis (PCA) indicated the two outlier samples in C4-2B and three outlier samples in 22RV1 data. After further quantifying anomalies with robust PCA methods (rPCA), those samples were removed from further analysis.

## Results

### High ADA is associated with aggressive prostate cancer

ADA levels were analyzed across different clinical samples to determine expression patterns in benign and cancerous prostate tissues. RNA *in situ* hybridization was done across five Tissue microarrays (TMA) also validated the finding that ADA was high in prostate tumors and levels were higher in high-grade tumors (N=345; benign (44), 3+3 (77), 3+4 (115), 4+3 (50), 8 and above (30))(p value<0.00001) (Fig 1A). RNA expression analysis of RNA-Seq data from TCGA (n=498) also showed that ADA transcript levels were increased in tumors with higher Gleason scores (N=498; 3+3=45, 3+4=146, 4+3=101, 8+ =206) (p value=0.00167) (Fig 1B). Immunohistochemical (IHC) staining of ADA protein was done in four TMAs with prostate biopsy tissue and matched benign cores from PCa patients. This analysis revealed that ADA was elevated in PCa tissues (p value= 0.0316) (Fig 1C). Kaplan-Meier survival analysis of the TCGA data set revealed that high ADA expression is also associated with poor progression-free survival (N: low ADA= 249, high ADA= 248) (p value=0.0057) (Fig 1D). The population was divided into low ADA and high ADA based on the median value of normalized ADA levels (RNA-Seq by Expectation Maximization, RSEM=5.249). As previously stated, ADA deaminates adenosine and converts it to inosine. Therefore, an increase in ADA activity will lead to increased conversion of adenosine to inosine, resulting in higher inosine and lower adenosine levels. In an independent set of urine samples from a PCa case-control study (benign, n= 98 and PCa, n=155), we found a high inosine-to-adenosine ratio (p value=0.0165) (Fig 1E) in PCa patients. The inosine-to-adenosine ratio was also found elevated in the urine samples of AA PCa patients compared to EA PCa patients (p value=0.0201) (Fig 1F). The Gleason score distribution among AA and EA PCa patients was as stated below. EA patients had 40 with Gleason 6, 22 with Gleason 7, and 13 with 8+ Gleason scores. AA patients had 40 with Gleason 6, 28 with Gleason 7, and 7 with 8+ Gleason scores. The other clinical characteristics of the samples are listed in Table 1.

**Fig 1.**
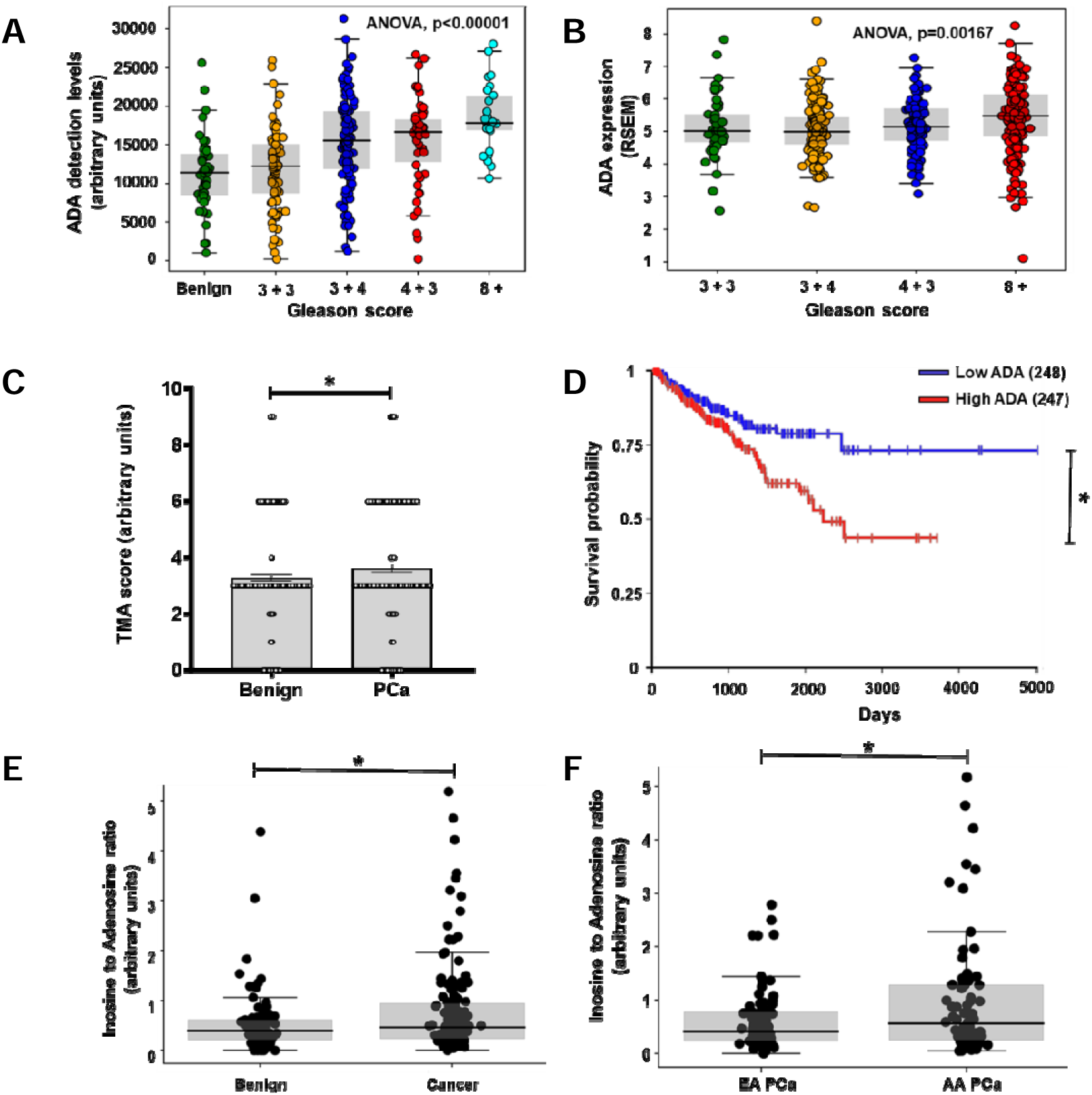
Elevated ADA in aggressive PCa. **A)** RNA scope analysis showed increased ADA levels in higher Gleason tumors (N: Benign= 3+3=77, 3+4=115, 4+3=50, 8+=30). **B)** ADA levels were found to be elevated with high Gleason scores in the TCGA prostate cancer dataset (N:3+3=45, 3+4=146, 4+3=101, 8+ =206).Normalized RNA sequencing data is represented as RSEM(RNA-Seq by Expectation Maximization) **C)** Immunostaining of TMAs for ADA showed that the ADA protein levels were high in PCa (N: Benign= 79, PCa= 77). **D)** Survival analysis done using the TCGA prostate cancer dataset revealed that high ADA in associated with poor survival in PCa (N: Low ADA-249, High ADA-248, Median= 5.249 RSEM) **E)** High inosine to adenosine ratio in the urine of PCa patients is indicative of high ADA enzyme activity (N: benign=98, cancer=155).The urine samples were obtained from a case-control study. **F)** High inosine to adenosine ratio in AA PCa compared to EA PCa (N: AA=78, EA=77) in the case-control cohort described in E. For comparison of ADA levels across Gleason scores, ANOVA, for Benign Vs Cancer comparison of ADA levels Wilcox matched-pairs signed rank test, for inosine to adenosine ratio in Benign Vs Cancer and EA Vs AA, Student’s unpaired two-tailed t-test were used to compute the p values. * p<0.05, ** p<0.01, *** p<0.001, **** p<0.0001

### Constitutive ADA overexpression decreases PCa cell adhesion

To understand the biological role of elevated ADA in PCa, ADA-overexpressing (ADA OE) cell line models were established using lentiviral transduction in MDA-PCa-2a (ancestry verified-AA cell line) and LNCaP (ancestry verified-EA cell line) cells. AA and EA cell lines were used to look for potential differences in outcomes between racial groups. To verify the specificity of the overexpression vector, a knockdown of ADA (ADA OE-KD) was performed on the ADA OE cell lines using shRNA (Fig 2). Therefore, ADA OE-KD cells contain both, ADA open reading frame (ORF) and shRNA and have intermediate expression levels of ADA. We expect ADA OE-KD to rescue (reverse) the effects observed in the ADA OE, thus ensuring that the changes observed are the consequence of ADA elevation and not any non-target effects. Vector control was established for both ADA OE (control) and ADA OE-KD (ADA OE-NT). The overexpression and knockdown models were verified by qPCR (mRNA), western blot (protein), ADA enzyme assay (enzyme activity), and LC/MS (adenosine and inosine levels) in both cell line models (Fig 2 A-H).

**Fig 2.**
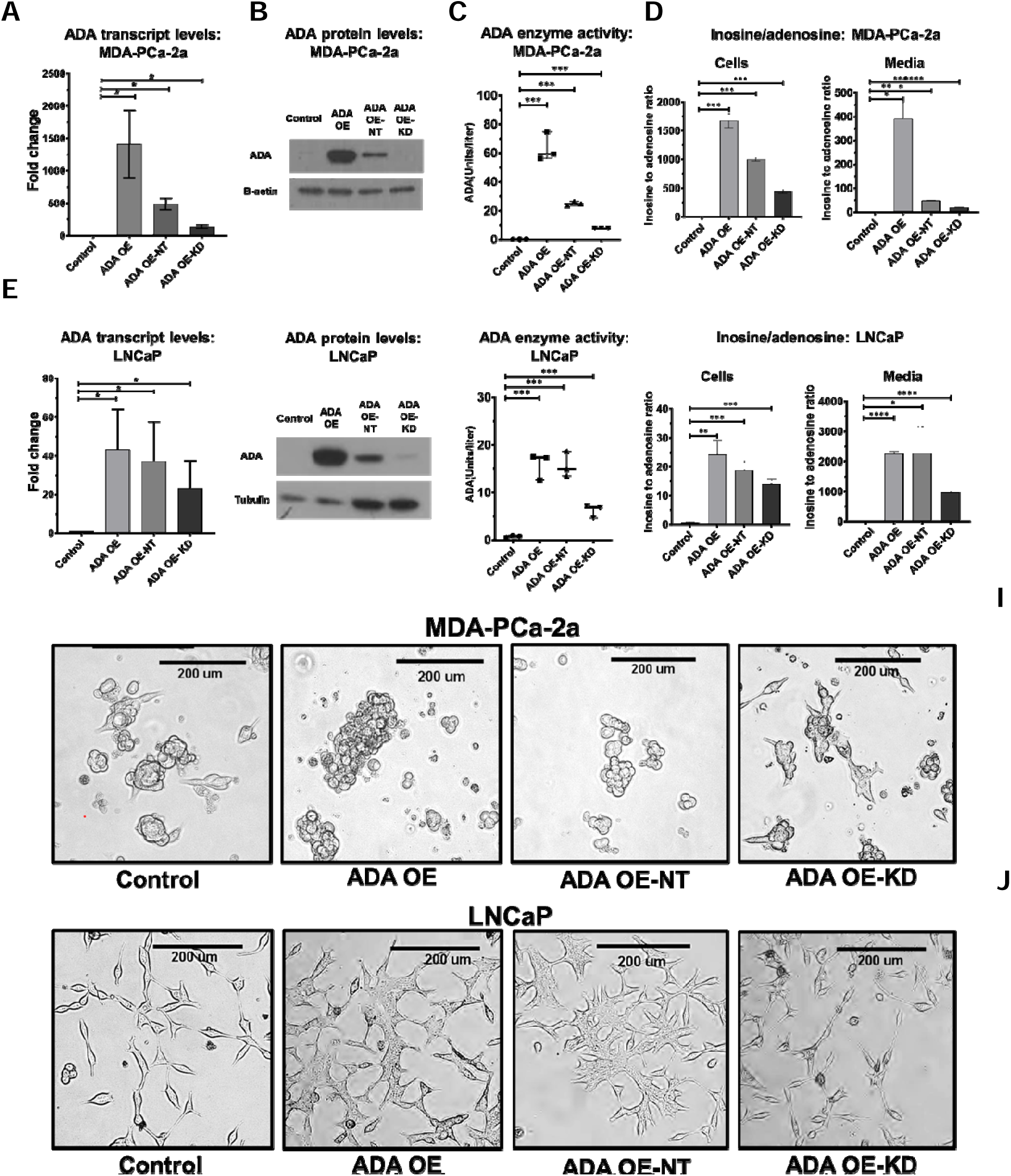
Establishing ADA OE in vitro models. Validation of established ADA OE and rescue models by qPCR, western blot, enzyme activity assay and LC/MS in **A-D)** MDA-PCa-2a (AA) and **E-H)** LNCaP cell lines(EA). Morphological changes in the **I)** MDA-PCa-2a and J) LNCaP cells. Student’s unpaired two-tailed t-tests was used for evaluating the statistical significance * p<0.05, ** p<0.01, *** p<0.001, **** p<0.0001

ADA OE cells exhibited morphological and adhesion changes compared to the control. This effect was also rescued in the ADA OE-KD cells (Fig 2 I&J). We used the Xcelligence real-time cell analysis system to quantify the changes in cell-surface adhesion. In both the cell lines, the cells with high ADA (ADA OE and ADA OE-NT) had decreased adhesion potential compared to the cells with lower ADA (control and ADA OE-KD) (Fig 3 A&B).

**Fig 3.**
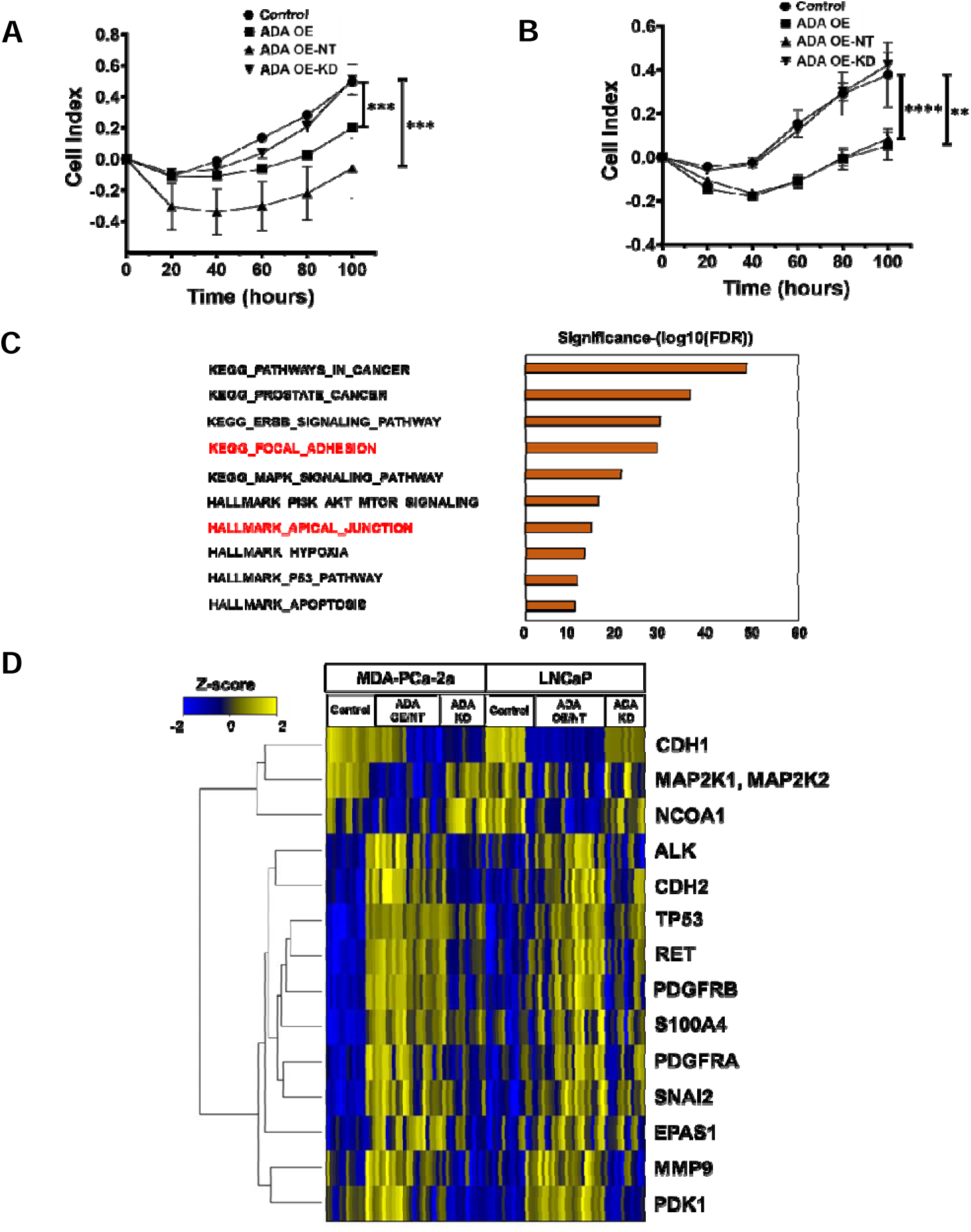
Constitutive ADA overexpression alters the cells’ adhesion potential. Decreased adhesion potential upon ADA OE observed in both **A)** MDA-PCa-2a and **B)** LNCaP cells (N=3/group/cell line). **C)** Gene set enrichment analysis (GSEA) of the RPPA data revealed that adhesion machinery was among the top altered mechanisms upon ADA elevation in both MDA-PCa-2a and LNCaP cells. Pathways altering cell adhesion are highlighted in red. **D)** RPPA analysis in both MDA-PCa-2A and LNCaP cells revealed alterations in several proteins. This heatmap shows proteins that are differentially expressed upon ADA OE (ADA OE and ADA OE-NT) in both MDA-PCa-2a and LNCaP cell lines (N=3/group/cell line). Heatmap is clustered by the log fold change in protein expression. Gradations of yellow and blue represent increased and reduced fold-change in the proteins, respectively. All GSEA concepts listed are significant at p<0.01 and FDR < 0.05. * p<0.05, ** p<0.01, *** p<0.001, **** p<0.0001

To further explore the molecular changes associated with this observed adhesion decrease, we utilized a reverse phase protein array (RPPA) platform for targeted proteomics analysis. 233 validated antibodies that include total and phosphoproteins of various pathways and functional protein groups were probed. Cells with high ADA levels showed changes in the cadherins and cadherin-regulating proteins (CDH1, CDH2, SNAI2), an increase in extracellular matrix (ECM) remodeling proteins (MMP9, S100A4), and several other proteins that are associated with invasion and metastasis (RET, PDFGRA, PDFGRB) (Fig 3D). Gene set enrichment analysis (GSEA) was conducted using KEGG and Hallmark pathways to identify significantly enriched pathways. We found that focal adhesion and apical junction pathways were two of the top enriched pathways upon ADA overexpression (Fig 3C).

### ADA overexpression delays tumorigenesis

In vivo, studies were done in NOD-SCID-Gamma (NSG) (MDA-PCa-2a, 22Rv1) and nude mice (for LNCaP only) to understand the role of ADA in PCa progression. The cells were injected subcutaneously, and their growth was monitored. It was evident that the presence of high ADA deterred the onset of tumors. Only the control tumors with baseline ADA levels had a successful take rate and developed tumors faster. This was consistent in all three cell lines used (MDA-PCa-2a, LNCaP, and 22Rv1). ADA OE cells formed tumors after 14 weeks (Vs 11 weeks in control, in MDA-PCa-2a (N=6 per group, p<0.001), 40 days (Vs 14 days in control) in LNCaP (N=8 per group, p<0.0001), and 14 days (Vs 9 days in control) in 22Rv1 cell lines (N=9 per group, p<0.0001) (Fig 4 A, B&C).

**Fig 4.**
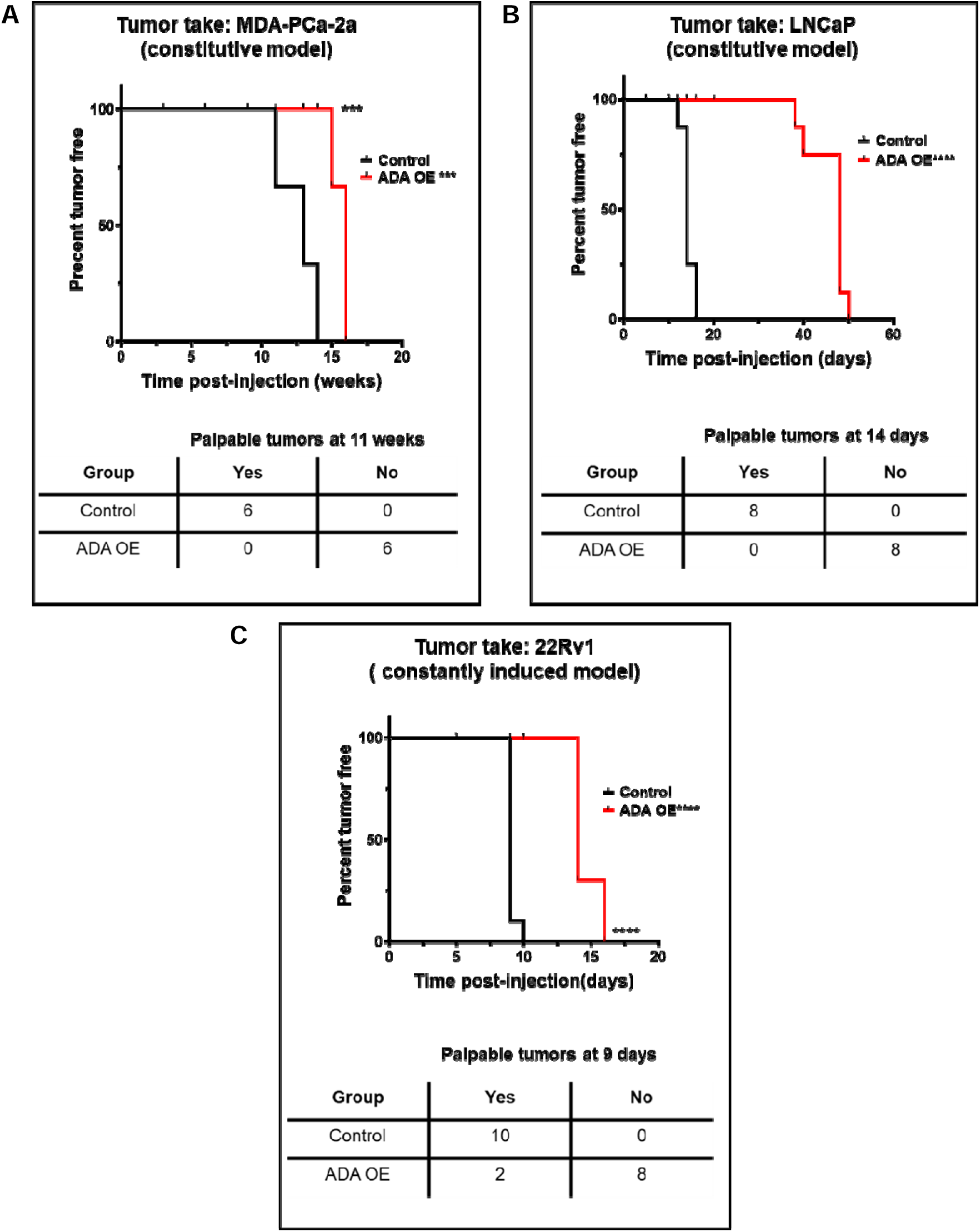
ADA OE delays tumor onset. Constitutive ADA expression delays the tumor onset in **A)** MDA-PCa-2a (N=6/group), **B)** LNCaP(N=8/group) and 22Rv1 (N=9/group) cells. Palpable tumors are typically ≤ 10 mm^3^. p-values were computed using log-rank test * p<0.05, ** p<0.01, *** p<0.001, **** p<0.0001

### Increased ADA expression enhances prostate tumor growth

To understand the significance of ADA at later stages in tumor progression, tet-inducible ADA-OE cell lines were used in vivo. 22Rv1 and C4-2B cell lines that are more robust and have better in vivo tumorigenicity were used for this study. The cells were injected without ADA induction and monitored until tumors were formed (∼50mm^3^). ADA expression was induced after the tumors reached 50mm^3^ and the growth rate was further monitored until the tumors reached 500mm^3^. Upon ADA induction, the tumors experienced a growth spurt. The ADA-OE-induced tumors grew faster than the control tumors in both 22Rv1 and C4-2B models (Fig 5 A&B). The tumor volumes were normalized to the volumes on the day of induction. The ratio of the final volume to the volume at the start of doxycycline induction was plotted as the outcome. The induction of ADA in the tumors was verified by qPCR and ADA enzyme activity assay (Fig 5 C&D).

**Fig 5.**
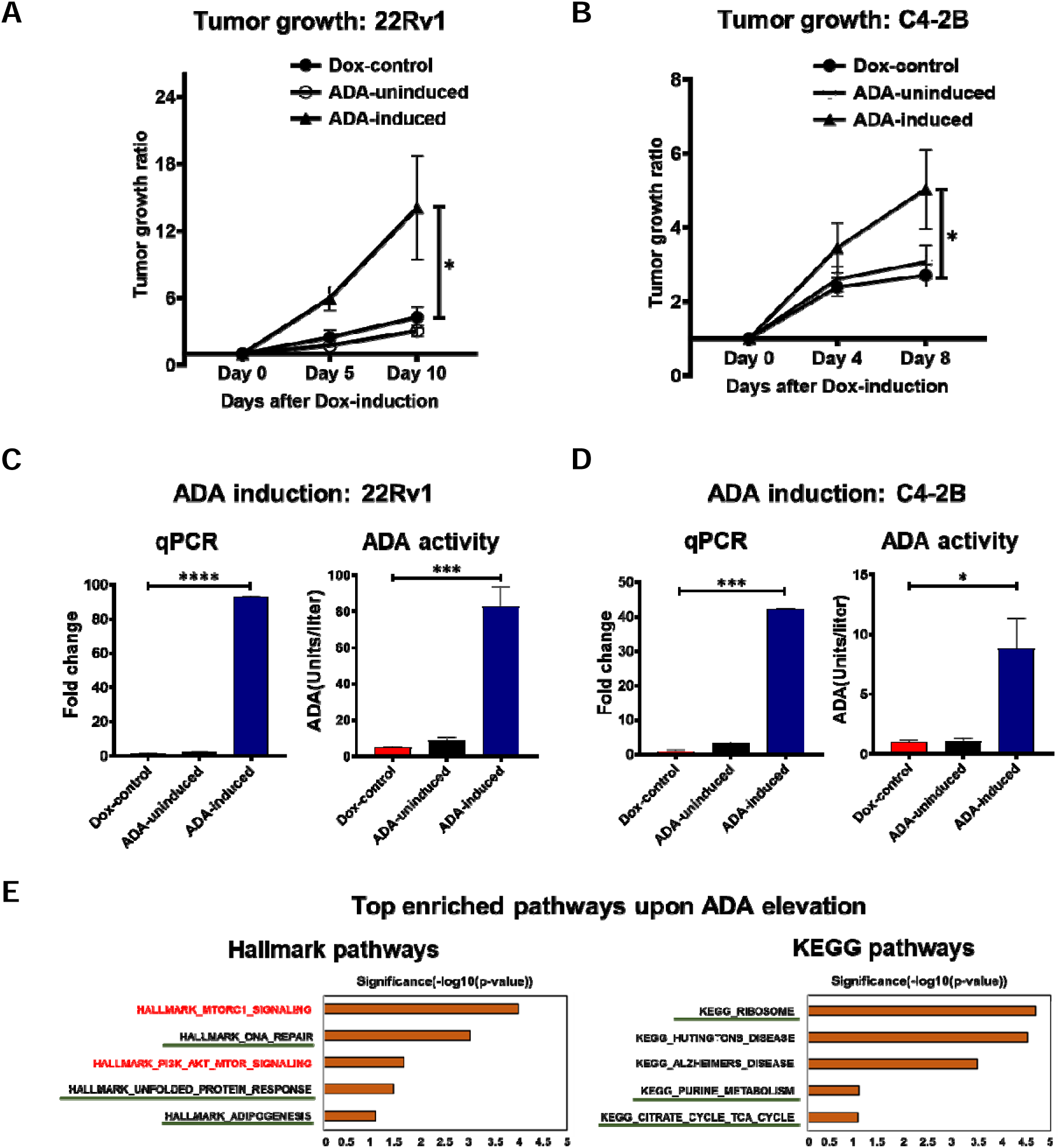
ADA elevation promotes prostate tumor growth. ADA elevation accelerates the tumor growth when induced post-tumor initiation in **A)** 22Rv1 (N: Dox-control=7, ADA-uninduced=7, ADA induced=8) and **B)** C4-2B (N: Dox-control=7, ADA-induced=9, ADA induced=8) ADA-inducible xenografts. Induction of ADA by doxycycline verified in by RT-qPCR and ADA enzyme assay in **C)** 22Rv1 and **D)** C4-2B tumors. **E)** GSEA of RNA sequencing data obtained from these tumors shows that mTOR signaling (highlighted in red) is enriched in tumors with high ADA. Pathways regulated downstream of mTOR were also enriched (underlined). All GSEA concepts listed are significant at p<0.05 and FDR < 0.25. p-vales were computed using Mann-Whitney test for tumor growth and Student’s two-tailed t-tests for ADA induction levels. * p<0.05, ** p<0.01, *** p<0.001, **** p<0.0001

Furthermore, RNA sequencing of the tumors followed by GSEA revealed that nutrient-sensing mTOR signaling was enriched upon ADA induction (Fig 5 E).

## Discussion

We study the metabolic landscape of PCa to understand the biochemical changes associated with PCa development and progression. We analyzed the clinical samples and identified that ADA was high in PCa. In addition, we also identified that ADA has a significant impact on tumor growth and cell adhesion.

Our clinical data analyses show that ADA levels are elevated in PCa, and correlated with the Gleason scores. Gleason scores ≤ 6 are considered to be indolent tumors and are considered for active surveillance. Gleason score 7, which consists of both 3+4 and 4+3 is considered intermediate grade with 4+3 being slightly worse than the 3+4 score. Scores ≥ 8 are considered highly aggressive, where progression and disease recurrence is highly likely [16, 17]. ADA levels being significantly high in higher Gleason scores indicates that the enzyme levels are associated with aggressive PCa.

High levels of ADA lead to increased conversion of adenosine to inosine. Therefore, under such conditions, the ratio of inosine to adenosine will be high. Our metabolic studies done on urine samples from a case-control study shows that the inosine-to-adenosine ratio is higher in PCa patients, especially in AA men with PCa who most likely have a poor clinical outcome [18, 19]. These findings further support the association of elevated ADA with aggressive PCa.

In vitro studies, conducted to delineate the role of ADA in PCa, revealed that when ADA is elevated for prolonged periods, the cells lose their ability to attach to the surface. Cells’ adhesion machinery maintains the organization and integrity of the cells in a tissue. It maintains the cells’ apical-basal polarity and is also important for regulating inter-cellular and extracellular communications. RPPA analysis revealed alterations in focal adhesion and apical junction pathways upon constitutive ADA overexpression. Focal adhesion pathway components maintain the connections between the cell’s cytoskeleton and extracellular matrix (ECM). It senses the changes in the microenvironment and alters the cell morphology and the cell’s adhesion to the ECM/basement membrane[20, 21]. The adhesion decrease observed in high ADA conditions could be a result of alterations in the focal adhesion pathway. The apical junction proteins are integral in maintaining cell-cell attachments and the polarity of the cells. In cancer, as the tumors progress and get dedifferentiated, the focal and apical junction complexes get altered significantly. The cells attain a more mesenchymal-like phenotype where they lose their polarity and adhesion potential. This makes them more motile and promotes migration and metastasis [21]. In the same RPPA study, we observed a decrease in E-cadherin levels and an increase in N-cadherin levels upon ADA OE. This cadherin switch is indicative of the epithelial-mesenchymal transition (EMT). Proteins S100A4 and SNAI2 which are upregulated in ADA OE cells are known to repress E-cadherins and some critical junction proteins, thus promoting the EMT phenotype [22, 23]. The adhesion capacity of the cells is also determined by ECM remodeling in addition to alterations in focal and apical complexes. S100A4 gets secreted into the extracellular space and facilitates ECM reconstitution by deploying matrix metalloproteases like MMP9 [23]. The adhesion decrease observed upon ADA elevation could be attributed to changes in these proteins and pathways. Adhesion decrease is significant in tumor progression during the later aggressive stages, especially during the time of metastatic dissemination. Several other proteins like RET, PDGFRA, and PDGFRB associated with invasion and metastasis[24–27] were also elevated upon ADA overexpression. These molecular alterations observed in vitro further support an association of ADA with aggressive PCa.

In vivo models are more relevant in determining the molecular pathophysiology of any disease. However, there are several challenges concerning in vivo models that are used to study PCa. It has been extremely hard to recapitulate the landscape of human prostate cancer in animal models. The existing prostate cell lines have a successful take rate only in immunodeficient mice and very rarely develop spontaneous metastasis. This poses us with the challenge of understanding the role of ADA in promoting metastasis and invasion PCa. Our in vivo tumorigenesis studies with the constitutive ADA OE cell lines lead to delayed tumor onset. This delay might be because of the altered adhesion that prevents them from attaching, thereby delaying the tumor take. While adhesion decrease is very critical in the later stages of tumor growth and progression, it can interfere with the process of tumor formation in mice models. This presented another challenge for us, improper tumor development from ADA OE cells prevented us from progressing to understand what happens in the later stages.

To circumvent the above challenges and to understand the role of ADA in promoting tumor growth, we generated inducible ADA OE models in more aggressive 22Rv1 and C4-2B cell lines. In the in vivo experiments, where ADA levels were regulated by doxycycline, there was a growth surge observed after the tumor onset. This suggests a role for elevated ADA in promoting growth during the later stages of PCa.

RNA sequencing analysis done on these tumors shed light on the potential mechanisms activated by ADA activity to promote tumor growth. The mTOR (mTORC1 and PI3K/AKT/mTOR) signaling was significantly enriched with a high enrichment score indicating its activation upon the increase in ADA activity. mTOR is a part of the mTORC protein complex. mTOR senses the cellular metabolic patterns and regulates the cell growth/survival signaling. Activation of mTOR promotes growth and survival by stimulating several biosynthetic pathways [28, 29]. An important role of mTOR is the promotion of protein synthesis by fostering ribosome biogenesis and thereby facilitating a higher translation rate. As a consequence of high protein production caused by mTOR activity, the unfolded protein response also is often upregulated [30–32]. mTOR also promotes cell growth by regulating mitochondrial functions, where it facilitates TCA anaplerosis by enhancing glutamine and glucose uptake. By elevating the TCA cycle, it provides substrates for purine synthesis which is vital transcription and energy homeostasis processes that ensure cells’ sustenance and survival [33–36]. The AKT-activated mTOR pathway is very critical for lipid biosynthesis and adipocyte development. mTORC1 is a well-established positive regulator of adipogenesis [37, 38]. Our analysis not only shows a positive enrichment of the mTOR signaling but also the downstream processes activated by mTOR like ribosome biogenesis, DNA repair, adipogenesis, purine synthesis, and TCA cycle are all enriched in tumors with elevated ADA. It has been well characterized that inosine elevation activates mTOR and the associated downstream metabolic biosynthetic and cell growth pathways. It is postulated that inosine activates mTOR via purinergic receptors [39, 40]. As described earlier, elevated ADA expression results in increased inosine levels which in turn could be responsible for activating the mTOR-driven pathways and promoting tumor growth in PCa.

Taken together our clinical, in vitro, and in vivo findings all point toward the significance of ADA upregulation in later aggressive stages of tumor. ADA enzymatic levels could experience a gradual surge as the tumor progresses. The initial surge after tumor formation could potentially lead to faster growth and progression by activating the mTOR signaling. While the initial elevation in ADA activity activates the cell growth and sustenance pathways, over time the prolonged/chronic ADA elevation could lead to alterations in the adhesion machinery and could potentially promote metastatic dissemination. The prolonged elevation allows the metabolites to alter the dynamics in the microenvironment and bring about changes in ECM and cell adhesion.

Our study for the first time has shown the intrinsic effects of ADA in any cancer with the use of extensive in vitro and in vivo models. We have shown evidence for ADA upregulation in PCa using clinical samples at transcript, protein, and metabolite levels. We have established both the acute and chronic effects of ADA upregulation on prostate tumors. This study further opens the prospects to evaluate the biomarker potential of inosine/adenosine, the mechanism associated with the alteration of growth/adhesion pathways by ADA, and the therapeutic potential of ADA in PCa in the future.

## Acknowledgments

The authors acknowledge the Global Center for Mass Spectrometry Excellence supported by Agilent Technologies at BCM. This research was partially supported by the following grants: Prostate Cancer Foundation Challenge Award (ASK, JAJ, and MI), Dan L. Duncan Cancer Center (P30 CA125123) supporting Human Tissue Acquisition and Pathology, and Metabolomics Shared Resources, CPRIT Metabolomics Core Facility (RP120092) supporting VP, NP and ASK, RO1CA227559 (Ask, GP), RO1CA267090 (ASK), and Translational Grant from V-Foundation (ASK).

